# Genome sequencing and molecular characterisation of XDR *Acinetobacter baumannii* reveal complexities in resistance: Novel combination of Sulbactam-Durlobactam holds promise for therapeutic intervention

**DOI:** 10.1101/2021.08.22.457246

**Authors:** Aniket Naha, Saranya Vijayakumar, Binesh Lal, Baby Abirami Shankar, Suriya Chandran, Sudha Ramaiah, Balaji Veeraraghavan, Anand Anbarasu

## Abstract

*Acinetobacter baumannii* is an emerging nosocomial strain expressing extensive drug resistance (XDR). Whole-genome sequencing and molecular characterisation analysis revealed the presence of carbapenemase in 92.86% of studied Indian isolates having *bla*_OXA-51_, *bla*_OXA-23_, *bla*_OXA-58_, and *bla*_NDM_ genes, with a few evidences of dual carbapenemase genes. As per the MLST scheme, IC2^Oxf^/CC2^Pas^ was the predominant clone, with 57.14% isolates belonging to this lineage. The presence of β-lactamases has rendered sulbactam (SUL) resistance (MIC: 16-256µg/ml) in all the studied isolates. The efficacy of novel durlobactam (DUR) in inhibiting β-lactamases and PBP2 was assessed through *in-silico* inter-molecular interaction analysis. Several non-synonymous single nucleotide polymorphisms (nsSNPs) were identified in PBP2 (G264S, I108V, S259T) and PBP3 (A515V, T526S) sequences. Minimal variations were recorded in the protein-backbone dynamics in active-site motifs of wild-type (WT) and mutants (MT), which correlated with the negligible binding energy fluctuations for PBP3-SUL (−5.85±0.04Kcal/mol) and PBP2-DUR (−5.16±0.66Kcal/mol) complexes. Furthermore, stronger binding affinities and low inhibition constants were noted in DUR complexed with OXA23 (−7.36Kcal/mol; 4.01µM), OXA58 (−6.44Kcal/mol; 19.07µM) and NDM (−6.82Kcal/mol; 10.01µM) when compared with conventional drugs avibactam and aztreonam. Stable interaction profiles of DUR, can possibly restore SUL activity against both PBP3_WT_ and PBP3_MTs_. The study establishes the efficacy of novel SUL-DUR combination as a successful treatment strategy to combat emerging XDR strains.

## INTRODUCTION

*Acinetobacter baumannii* (*A. baumannii*), a non-fermenting Gram-negative coccobacillus is responsible for several nosocomial outbreaks causing bloodstream infections, ventilator-associated pneumonia, urinary-tract infections and wound infections [Bassetti et al., 2021; Sagan et al., 2020]. There has been a growing concern among the clinicians in treating multi-drug resistant (MDR) strains of *A. baumannii* infections [Bassetti and Giacobbe, 2020]. Recently, the WHO categorised carbapenem-resistant *A. baumannii* (CR*Ab*) under priority-1: critical pathogen for research and development of new antibiotics (www.who.int). In combination with other antibiotics, colistin is currently administered for treating CR*Ab* strains. Excessive colistin usage resulted in increased antimicrobial resistance (AMR) and colistin-induced adverse reactions [Rodriguez et al., 2020]. Minocycline and tigecycline shows good *in-vitro* activity [Piperaki et al., 2019] while cefiderocol and eravacycline, are used as salvage therapy against CR*Ab* and could potentially become first-line choices for treating *A. baumannii* infections [Portsmouth et al., 2018; Solomkin et al., 2017]. However, newer β-lactams and β-lactamase inhibitors (βL-βLIs) combinations like ceftolozane-tazobactam, ceftazidime-avibactam, imipenem-relebactam and meropenem-vaborbactam are clinically ineffective against *A. baumannii* [Bassetti and Giacobbe, 2020; Tooke et al., 2019].

Recently, sulbactam-based regimens are growing interest and could be a possible alternative therapy. Sulbactam (SUL), though a β-lactamase inhibitor has intrinsic antibacterial affinity towards PBP3 and PBP1 of *A. baumannii* and therefore, considered a β-lactam antibiotic [Bassetti and Giacobbe, 2020; Yahav et al., 2020]. However, presence of various β-lactamases could limit its use [Butler et al., 2019]. Durlobactam (DUR), a non-βL-βLI has diazabicyclooctane (DBO) scaffold which inhibits a wide range of β-lactamases (primary target) and PBP2 (secondary target) [Yahav et al., 2020]. Recent studies demonstrated the potential antibacterial activity of the novel SUL-DUR (βL-βLI) inhibitor combination which is currently under phase-III clinical trials and might be helpful in treating infections caused by *A. baumannii* [McLeod et al., 2020; Yahav et al., 2020]. Our research group has worked extensively to understand the protein-ligand interaction and complex AMR mechanisms of pathogenic bacteria through genomic analyses [Jacob et al., 2019; Vasudevan et al., 2020; Vijayakumar et al., 2016, 2020], *in-silico* structural [Basu et al., 2020, 2021; Lavanya et al., 2015; Thillainayagam et al., 2020], and systems biology approaches [Debroy et al., 2020; Miryala et al., 2020, 2021; Naha et al., 2020]. The aim of the present study was to decipher the activity of this βL-βLI combination against *A. baumannii* by characterising the inter-molecular binding profiles of SUL with PBP3 and the potency of DUR to restore the efficacy of SUL by inhibiting the β-lactamases and PBP2 through *in-silico* and *in-vitro* analyses.

## MATERIALS AND METHODS

### Bacterial isolates

A total of 28 clinical strains of *A. baumannii* were identified from routine blood (n=21) and sputum cultures (n=7) from 2018-19 at Christian Medical College, Vellore. All the isolates were characterised up to the species level as *Acinetobacter baumannii calcoaceticus* complex (*Acb* complex) using routine biochemicals and automated identification system using VITEK^®^-MS (BioMérieux) [Vijayakumar et al., 2016].

### Antimicrobial susceptibility testing (AST) and Minimum inhibitory concentration (MIC)

AST was performed for all the isolates against 18 different antibiotics by the Kirby-Bauer disc-diffusion (DD) method and interpreted according to Clinical Laboratory Standards Institute (CLSI) guidelines [Clinical and Laboratory Standards Institute, 2021]. MIC for colistin and SUL were determined for all the study isolates using broth micro-dilution (BMD) and interpreted accordingly. Two quality control (QC) strains, *Escherichia coli* ATCC 25922 and *Pseudomonas aeruginosa* ATCC 27853 were used for colistin-MIC. As an internal control, *mcr-1* positive *E. coli* strain was also included. Additionally, two in-house QC strains, *Klebsiella pneumoniae* BA38416 (MIC 0.5µg/ml) and *Klebsiella pneumoniae* BA25425 (MIC 16µg/ml), were also included in every batch of testing. For SUL-MIC, *A. baumannii* NCTC 13304 was used as QC strain [Clinical and Laboratory Standards Institute, 2021].

### Molecular characterisation of carbapenemase genes by multiplex PCR

All 28 isolates were grown on blood-agar overnight, and genomic DNA was extracted using QIAamp DNA Mini Kit (QIAGEN, Hilden, Germany) as per the manufacturer’s instructions. Conventional multiplex-PCR was done to detect carbapenemase genes such as *bla*_KPC_, *bla*_IMP_, *bla*_VIM_, *bla*_NDM_, *bla*_SIM_, *bla*_OXA-48_, *bla*_OXA-23_, *bla*_OXA-24_ and *bla*_OXA-58_ like as described previously [Vijayakumar et al., 2020]. The amplicons were visualised in 2% agarose gel stained with ethidium-bromide. Known positive controls for appropriate genes were used in all the runs. (Courtesy: IHMA, Inc., USA).

### Whole-genome sequencing (WGS), assembly and annotation

WGS was performed with the extracted DNA for all 28 isolates. The quantity and quality of the DNA were analysed using the Qubit 3.0 fluorometer (Thermo Fisher, USA) and the Nanodrop spectrophotometer (Thermo Fisher, USA). In brief, short-read sequencing was performed using IonTorrent™ Personal Genome Machine™ (Life Technologies, Carlsbad, CA) with 400-bp read chemistry or paired-end genomic libraries were prepared and sequenced using Illumina HiSeq V4 platform as per the manufacturer’s instructions. Long-read sequencing was performed using Oxford Nanopore Technologies, Oxford, UK using 1D sequencing method according to the manufacturer’s protocol. To obtain complete genome, hybrid assembly was performed as described previously [Vasudevan et al., 2020].

The NCBI Prokaryotic Genome Annotation Pipeline was used to assemble and annotate all the genomes. Further downstream analysis was performed using CGE server (http://www.genomicepidemiology.org/) where genomes were analysed for the presence of acquired AMR genes using ResFinder v4.1 (https://cge.cbs.dtu.dk/services/ResFinder/) [Bortolaia et al., 2020]. The sequence type of the isolates was assigned by the MLST v2.0 tool using both the Oxford and Pasteur scheme (https://cge.cbs.dtu.dk//services/MLST/). A maximum-likelihood (ML) tree was constructed using RAxML v0.7.4 [Kozlov et al., 2019] with 28 whole-genomes, and visualised using iTOL v6.0 (https://itol.embl.de/) [Letunic and Bork, 2019]. Each genome was annotated with susceptibility profile of carbapenem SUL and colistin, International Clones and chromosomal mutation profile.

### Nucleotide accession numbers

The complete genome project of 28 studied strains were deposited at DDBJ/ENA/GenBank under accession numbers SP304-(**CP040080**), VB23193-(**CP035672**), VB473-(**CP050388**), VB958-(**CP040040**), VB1190-(**CP040047**), VB16141-(**CP040050**), VB31459-(**CP035930**), CIAT758-(**CP038500**), ACN21-(**CP038644**), VB35179-(**CP040053**), VB35435-(**CP040056**), VB33071-(**CP040084**), VB35575-(**CP040087**), VB2200-(**CP050421**), VB2486-(**CP050403**), P7774-(**CP040259**), VB82-(**CP050385**), VB7036-(**CP050523**), VB11737-(**CP050400**), VB723-(**CP050390**), PM2235-(**CP050410**), PM2696-(**CP050412**), PM3665-(**CP050415**), PM4122-(**CP050425**), PM4229-(**CP050432**), VB2107-(**CP051474**), VB2139-(**CP050526**) and VB2181-(**CP050401**).

### Mutation Analysis of Penicillin Binding Proteins (PBPs) and β-lactamases (βLs)

The PBP2, PBP3, OXA23, OXA58, and NDM sequences extracted from whole-genomes were scanned for non-synonymous single nucleotide polymorphisms (nsSNPs) using SeaView5 [Gouy et al., 2020] and were compared with the reference *A. baumannii* ATCC17978 (**CP000521**). Upon nsSNP detection, the stability at structural-functional aspects of the mutations was assessed with DUET server (http://structure.bioc.cam.ac.uk/duet). Thermodynamic free-energy changes (ΔΔG) are computed upon site-directed mutagenesis leading to alteration in protein stability [Pires et al., 2014]. Further, the destabilisation effects of nsSNPs were estimated by N-H S^2^ bond-order using DynaMine server (http://dynamine.ibsquare.be/). The movement restrictions of atomic bond-vectors (S^2^-scores) are derived directly from experimentally validated NMR chemical shifts [Cilia et al., 2014].

### *In-silico* Protein Modelling

NCBI-BLASTp analysis against RCSB-PDB retrieved suitable templates for PBP3-3UE3, OXA23-5WI3, OXA58-4OH0 and NDM-4EY2. Global-alignment (https://www.ebi.ac.uk/Tools/psa/emboss_needle/) revealed 76.1% identity of PBP3 with 3UE3. Other targets shared high identities of 98.4%, 99.6%, and 100% with OXA23, OXA58, and NDM sequences, respectively. Lack of templates for PBP2 and low (<90%) sequence identity for PBP3 appealed *in-silico* protein modelling. An extended dual-step protein modelling was performed as reported previously [Basu et al., 2021].

### Protein Structure Refinement and Validation

The modelled proteins (PBP2, PBP3) and β-lactamases (OXA23, OXA58, and NDM) were scrutinised for several poor-rotamers, unstable bond-torsions and outliers. These proteins were refined using GalaxyRefine server (http://galaxy.seoklab.org/index.html) to enhance the structure quality by furnishing and repacking the side-chains [Heo et al., 2013]. Energy-minimization was performed using Swiss-PDB-Viewer v4.1.0 *in-vacou* with GROMOS96-43B1 Force-Field by carrying out 2000 steps each for steepest-descent and conjugate-gradients [Kaplan and Littlejohn, 2001]. The nsSNPs were introduced in wild-type (WT) protein-targets to generate respective mutants (MT) using SPDBV. The secondary protein structures were assessed using SOPMA (https://npsa-prabi.ibcp.fr/cgi-bin/npsa_automat.pl?page=npsa_sopma.html) and tertiary structures were validated with ProSA-web [Wiederstein and Sippl, 2007], HARMONY [Pugalenthi et al., 2006], and ProTSAV [Singh et al., 2016] servers. UCSF-CHIMERA was used for protein structure visualisation [Pettersen et al., 2004].

### Active-site prediction

Several reports highlighted three conserved motifs (S-X-X-K, S-X-N/D and K-T/S-G-T) within PBPs playing pivotal roles in βL binding [King et al., 2017; Rajavel et al., 2021]. The S-X-X-K motif in β-lactamases was conserved with PBP2 [Adachi et al., 1992] while active-sites of NDM were retrieved from literary evidences [Yuan et al., 2012] and validated through CASTp-3.0 server (http://sts.bioe.uic.edu/castp/calculation.html) [Tian et al., 2018].

### Ligand selection

The ligands for the present study were procured from PubChem Database (https://pubchem.ncbi.nlm.nih.gov/). The 3D-coordinates of SUL (130313), DUR (89851852), Avibactam (AVI) (9835049), and Aztreonam (AZT) (5742832) were generated using CORINA server (https://www.mn-am.com/online_demos/corina_demo). The polar H-atoms were added and spatial arrangements of ligands were visualised in Avogadro software v1.2.0 [Hanwell et al., 2012].

### Molecular Docking Analysis

Refined proteins were subjected to Molecular Docking Analysis with AutoDock 4.2. Both the WT and MT protein-targets were optimised prior to docking analysis by adding polar H-atoms, Kollman charges and merging non-polar H-atoms stabilizing the free-end residues of the protein moiety. The ligand torsions were fixed upon the addition of Gasteiger charges. The affinity grid box was constructed centering the active-site residue coordinates of dimensions 60Å^3^ with equal spacing of 0.375Å. Docked-poses were generated based on Lamarckian and genetic algorithms, while protein-ligand complex with the least binding-energies (BE) were selected for the present study [Morris et al., 2009]. The docked complex depicting several inter-molecular interactions were visualised in Discovery Studio 2020 v20.1.0.19295 [Jejurikar and Rohane, 2021]. Molecular Docking of SUL was performed with PBP3 while DUR was docked with β-lactamases and PBP2. The classical drugs AVI and AZT were used as positive controls against both DUR-β-lactamase and DUR-NDM complexes respectively.

## RESULTS AND DISCUSSION

### Bacterial isolates and antimicrobial susceptibility

All 28 *Acb* complexes were confirmed as *A. baumannii* using VITEK^®^-MS and re-confirmed by the presence of an intrinsic *bla*_OXA-51_ like gene by uniplex-PCR. AST revealed six isolates as PDR being resistant to all the tested antibiotics while 21 showed XDR profiles being resistant to all the tested antibiotics except for minocycline and tigecycline (Blood, n=20 and sputum, n=6). Of the remaining two isolates, one from blood was MDR (PM4229) while another isolate (SP304) from sputum was PDS [**Supplementary File S1**]. All isolates were SUL resistant with four from blood and one from sputum were colistin intermediate respectively, while the remaining 23 isolates were colistin susceptible (Blood, n=17 and sputum, n=6). The SUL and colistin MIC range, MIC_50_ and MIC_90_ of the blood and sputum isolates were tabulated [**Table 1**].

**Table 1:**
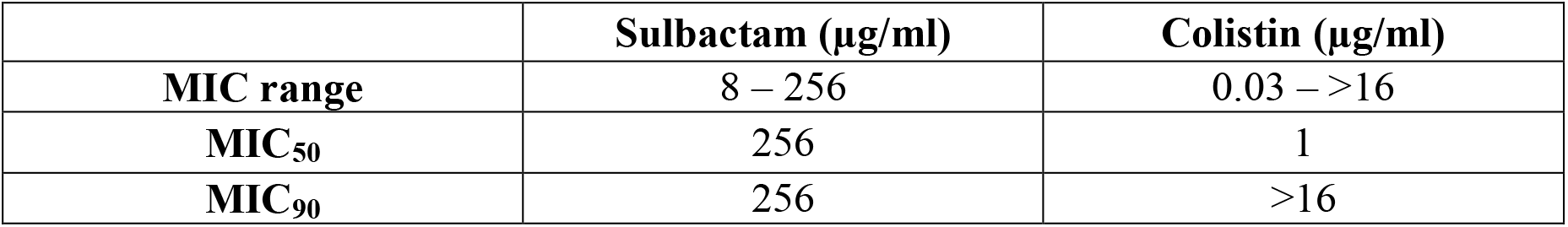
MIC range, MIC_50_ and MIC_90_ of clinical strains of *A. baumannii* tested against Sulbactam and Colistin (n=28)

|  | Sulbactam (µg/ml) | Colistin (µg/ml) |
| --- | --- | --- |
| **MIC range** | 8 – 256 | 0.03 – >16 |
| **MIC<sub>50</sub>** | 256 | 1 |
| **MIC<sub>90</sub>** | 256 | >16 |

### Carbapenemase PCR and WGS

Multiplex-PCR for carbapenemase genes showed the predominance of class-B and D carbapenemase, *bla*_OXA-23_ and *bla*_NDM_ like genes [**Figure 1**]. Only two isolates were positive for *bla*_OXA-58_ like. WGS revealed eight variants of intrinsic *bla*_OXA-51_ gene (*bla*_OXA-66_, *bla*_OXA-64,_ *bla*_OXA-68,_ *bla*_OXA-94,_ *bla*_OXA-144,_ *bla*_OXA-203,_ *bla*_OXA-337_ and *bla*_OXA-371_), two variants of *bla*_OXA-23_ (*bla*_OXA-23_ and *bla*_OXA-169_), two variants of *bla*_OXA-58_ (*bla*_OXA-58_ and *bla*_OXA-420_) and a single variant of *bla*_NDM_ (*bla*_NDM-1_). Fourteen isolates carried *bla*_OXA-23_ alone, ten isolates co-harboured *bla*_OXA-23_ and *bla*_NDM-1_, one was co-producer of *bla*_OXA-58_ and *bla*_OXA-23_ while another co-carried *bla*_OXA-420_ and *bla*_NDM-1_. The remaining two isolates were identified as carbapenemase negative. Based on the Oxford and Pasteur’s MLST schemes, 28 isolates were categorised into four International Clonal lineages (ICs) and one Clonal Complex (CC). Four novel STs were identified, while IC2^Oxf^/CC2^Pas^ was the predominant clone identified in 16 isolates, followed by IC8^Oxf^/CC10^Pas^ in three, IC7^Oxf^/CC25^Pas^ in two, IC1^Oxf^/CC1^Pas^ and CC85^Oxf^ each in one isolate. The ML phylogenetic tree from 28 whole-genomes was mapped to the reference genome *A. baumannii* ATCC 17978 (**CP012004**) [**Figure 2**].

**Figure 1:**
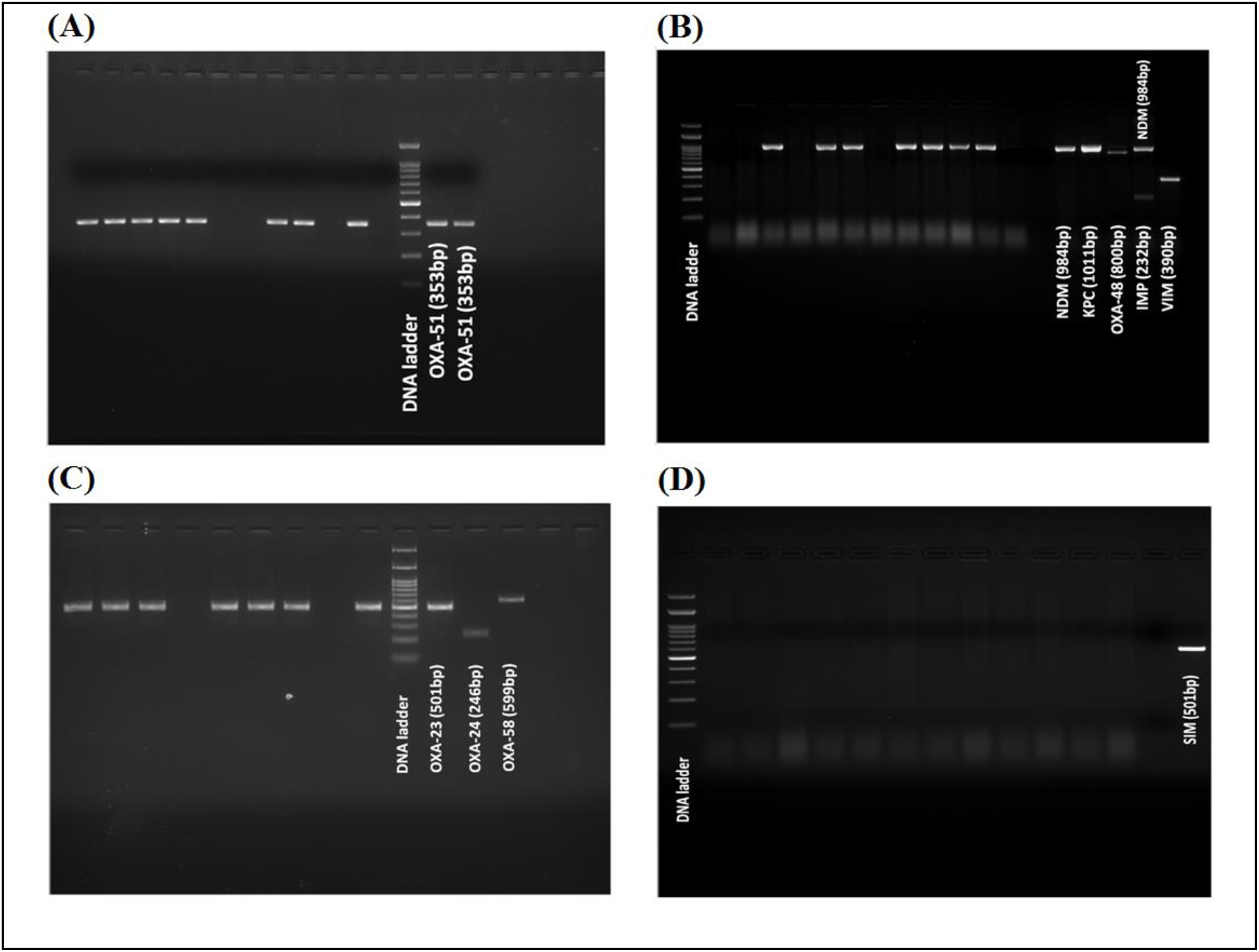
Gel electrophoresis of carbapenem resistant genes in *A. baumannii*: **(A)** Intrinsic *bla*_OXA-51_ gene **(B)** Different class of carbapenem resistant genes **(C)** Class D *bla*_OXA-23_, *bla*_OXA-24_, and *bla*_OXA-58_ genes **(D)** Class B *bla*_SIM_ gene

**Figure 2:**
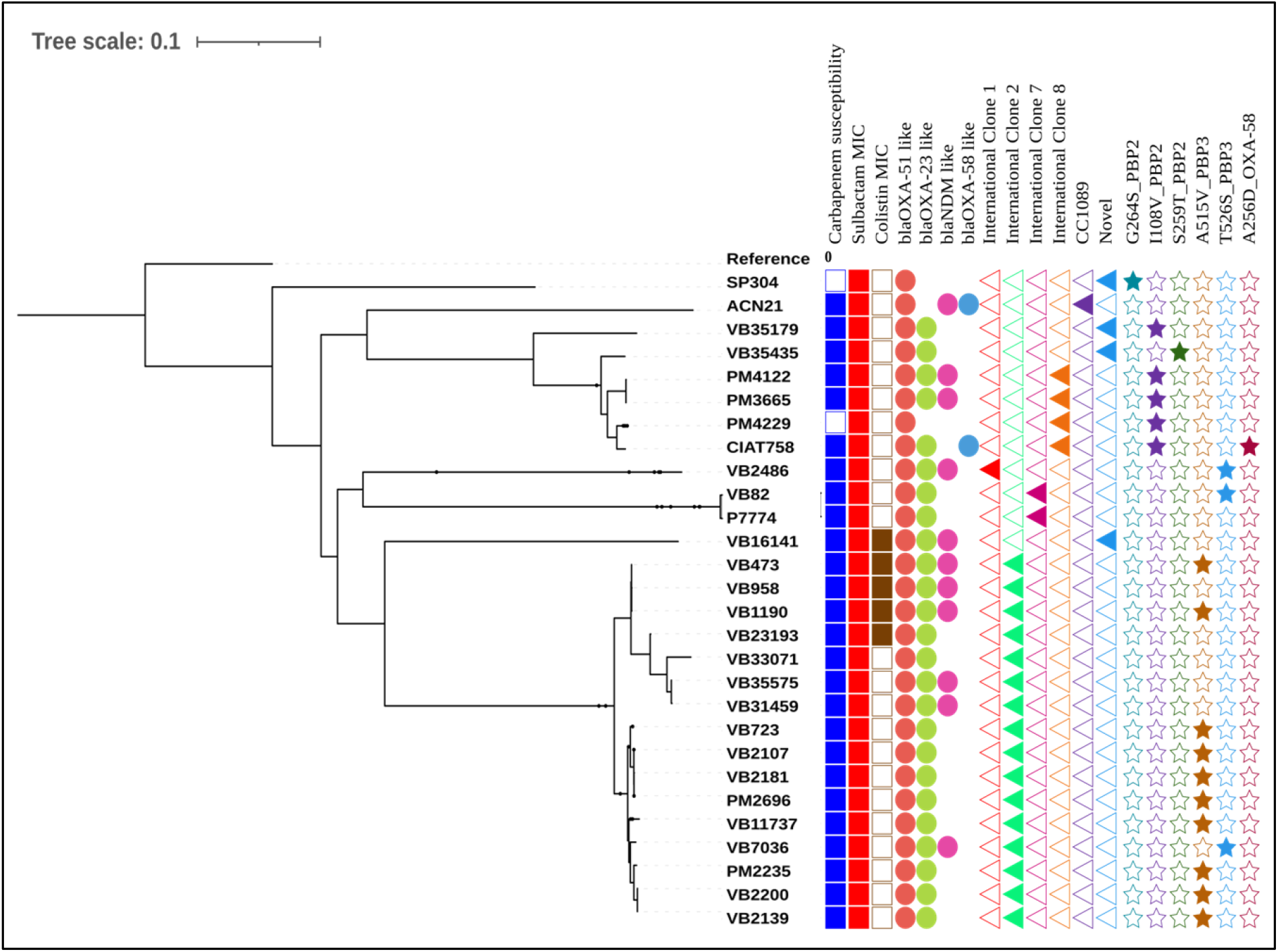
A maximum-likelihood phylogeny tree of 28 whole-genomes sequenced in this study. The colour filled boxes under carbapenem and sulbactam MIC indicated resistant, while for colistin MIC indicated intermediate and the empty boxes indicated susceptible. The colour filled shape denoted presence while the empty shape denoted absence of the respective traits

### Screening of nsSNPs from target proteins

PBP2_I108V_ (17.85%) was the most prevalent mutant allele while PBP2_G264S_ and PBP2_S259T_ were found in one isolate each. Amongst PBP3_MTs_, PBP3_A515V_ (35.71%) was predominant than PBP3_T526S_ (10.71%) mutant allele. *In-vitro* assay revealed increased SUL-MIC (16-256µg/ml) with either of these PBP3_MTs_. Three isolates (VB473, VB1190, and VB7036) with dual carbapenemase (*bla*_OXA-23_, *bla*_NDM-1_) genes had MIC of 256µg/ml. All PBP3_A515V_ isolates belonged to IC2^Oxf^/CC2^Pas^ clone, while 80% PBP2_I108V_ isolates belonged to IC8^Oxf^/CC10^Pas^ lineage. No mutations were detected in OXA23 and NDM, whereas the CIAT758 strain possessed A256D mutation in OXA58. The OXA23 and NDM were flanked with insertion sequence (IS) element IS*Aba1* (Tn*2006*) and IS*Aba125* (Tn*125*), respectively. These composite transposons attributed in their easy spread and efficient transmission, which resulted in the high prevalence of OXA23 and NDM in the isolates. Though IS*Aba3* flanks OXA58, the presence of transposon has not been reported so far, thereby lowering its prevalence and spread [Pagano et al., 2016]. The backbone dynamics of the neighbouring residues (amino-acid: 239-272) of OXA58_A256D_ revealed a 0.83% reduction in backbone rigidity compared with the OXA58_WT_. Owing to its flexible backbone dynamics, the enzymatic degradation capability of OXA58_MT_ is thus enhanced which might be a promising cause of gaining A256D mutation in OXA58.

The negative ΔΔG values from DUET servers signified the destabilising effects of nsSNPs at structural-functional levels. The extent of destabilisation was further validated with N-H S^2^ bond-order restraints [**Supplementary File S2(A)**]. Residue-level analysis of PBP2_I108V_ (S^2^-score=0.88) and PBP2_S259T_ (S^2^-score=0.9) revealed rigid-backbone conformers while PBP2_G264S_ had relatively flexible-backbone (S^2^-score=0.78) when compared with PBP2_WT_. The PBP2_G264S_ mutation (ΔΔG=-1.2Kcal/mol) had the lowest ΔΔG scores amongst PBP2_MTs_ and therefore, showed destabilisation profiles leading to protein unfolding at low energy enforcements. Similarly, PBP2_I108V_ (−1.123Kcal/mol) and PBP2_S259T_ (−0.476Kcal/mol) correlated with the most stable backbone dynamics. The PBP3_T526S_ mutation resulted in protein flexibility (S^2^-score=0.63), while reduced flexibility (S^2^-score=0.8) was observed in PBP3_A515V_ mutation. Thus, the backbone stability profiles established the lower energy requirement for protein unfolding upon PBP3_T526S_ (−1.45Kcal/mol) than PBP3_A515V_(0.34Kcal/mol) mutation. Thus, the protein-backbone dynamics validated the free-energy changes upon mutations and their effects on protein stability.

### *In-silico* structure prediction, refinement and validation

The protein-targets were refined and subjected to multiple validations. The validated modelled structure of PBP2 and PBP3 were submitted to Protein Model Database (PMDB) (http://srv00.recas.ba.infn.it/PMDB/), having PMDB-IDs PM0084111 and PM0084112, respectively. Lower MolProbity scores, minimum percentages of poor-rotamers and Ramachandran favoured zones (>95%) defined stable protein conformation with reduced steric-clashes and outliers. The protein structures and the active-sites predominantly constituted of random coils. The proteins had solvent accessibility area of 76.86±1.4% while 23.94±6.8% was buried [**Supplementary File S3**]. Global Z-scores signified that all target-proteins were energetically stable as they fell within the defined zones of experimentally characterised proteins **[Figure 3(A)]**. HARMONY propensity-calibration plots revealed that all the target-proteins lay extremely close to straight-fit line revealing negligible gross-errors in folding patterns [**Figure 3(B)**]. The ProTSAV heat-maps revealed unified Z-score, confirming protein structure quality within 2-5Å RMSD, making them ideal for protein-ligand interaction studies [Singh et al., 2016] [**Figure 3(C)**].

**Figure 3:**
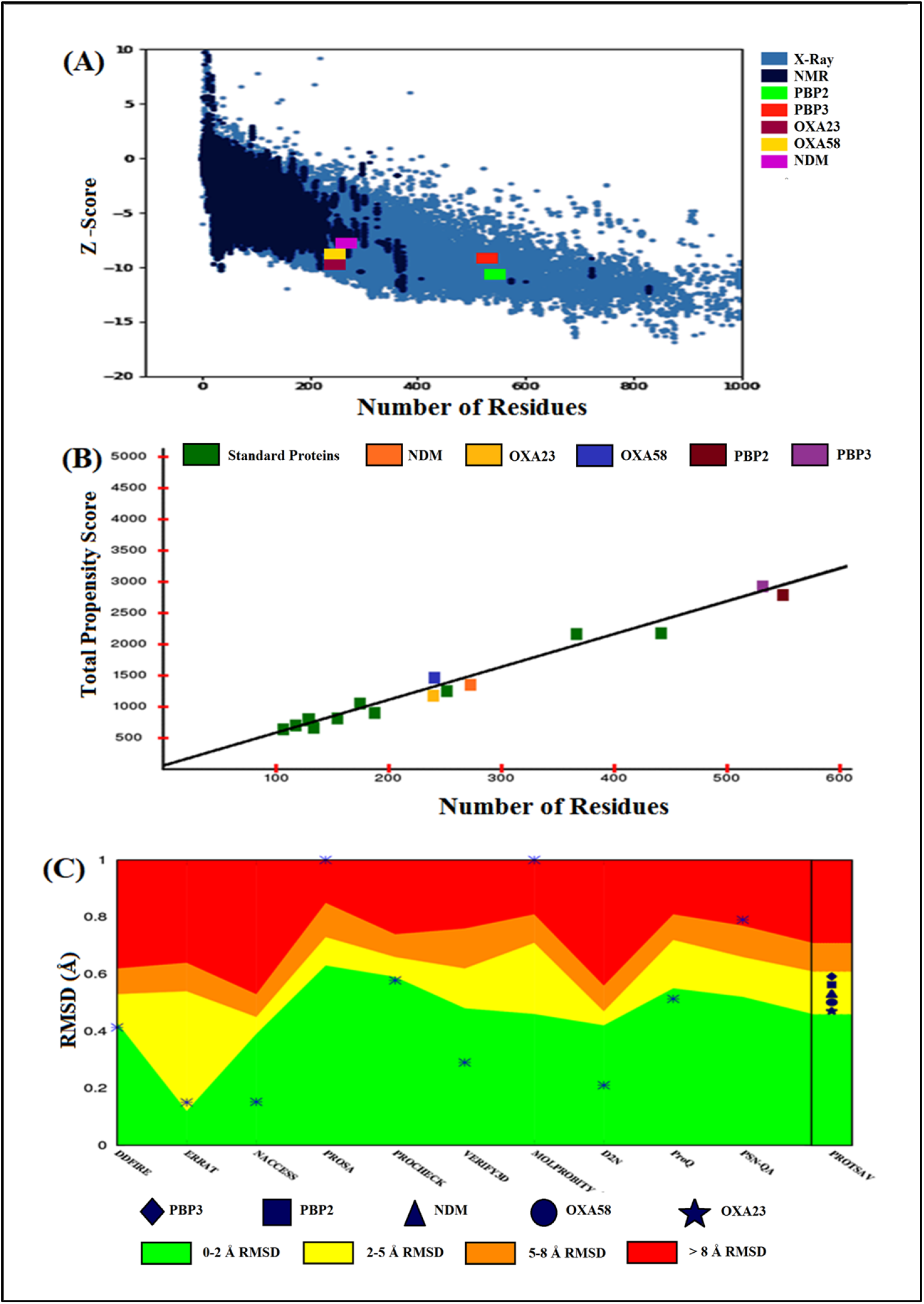
Structure evaluations of modelled proteins: **(A)** Global model quality of protein-targets **(B)** Propensity plots of protein-targets **(C)** ProTSAV heat-maps of protein-targets

### Active-site validations in target-proteins

The three conserved motifs S-X-X-K, S-X-N/D, and K-T/S-G-T were present in both PBP2 and PBP3 as [S326-T327-I328-K329] and [S336-T337-M338-K339]; [S383-C384-D385] and [S390-S391-N392]; [K537-T538-G539-T540] and [K525-T526-G527-T528] respectively. The S-X-X-K motif for OXA23 [S79-T80-F81-K82] and OXA58 [S83-T84-F85-K86] were retrieved along with other key residues of OXA23 (F110, M221) and OXA58 (F114, M225). In NDM, N220, V73, and D124 residues formed stable interactions with the ligands. The active-site motifs/residues of all target-proteins were validated with CASTp server prior to molecular docking analysis.

### Molecular Docking of PBPs and β-lactamases with βL-βLI combination

The variation in BEs of SUL-PBP3 and DUR-PBP2 complexes were analysed and correlated with the N-H S^2^ bond-order restraints to decipher the effect of nsSNPs on the binding profiles of the complexes. The BEs of protein-ligand complexes are represented in **Figure 4**.

**Figure 4:**
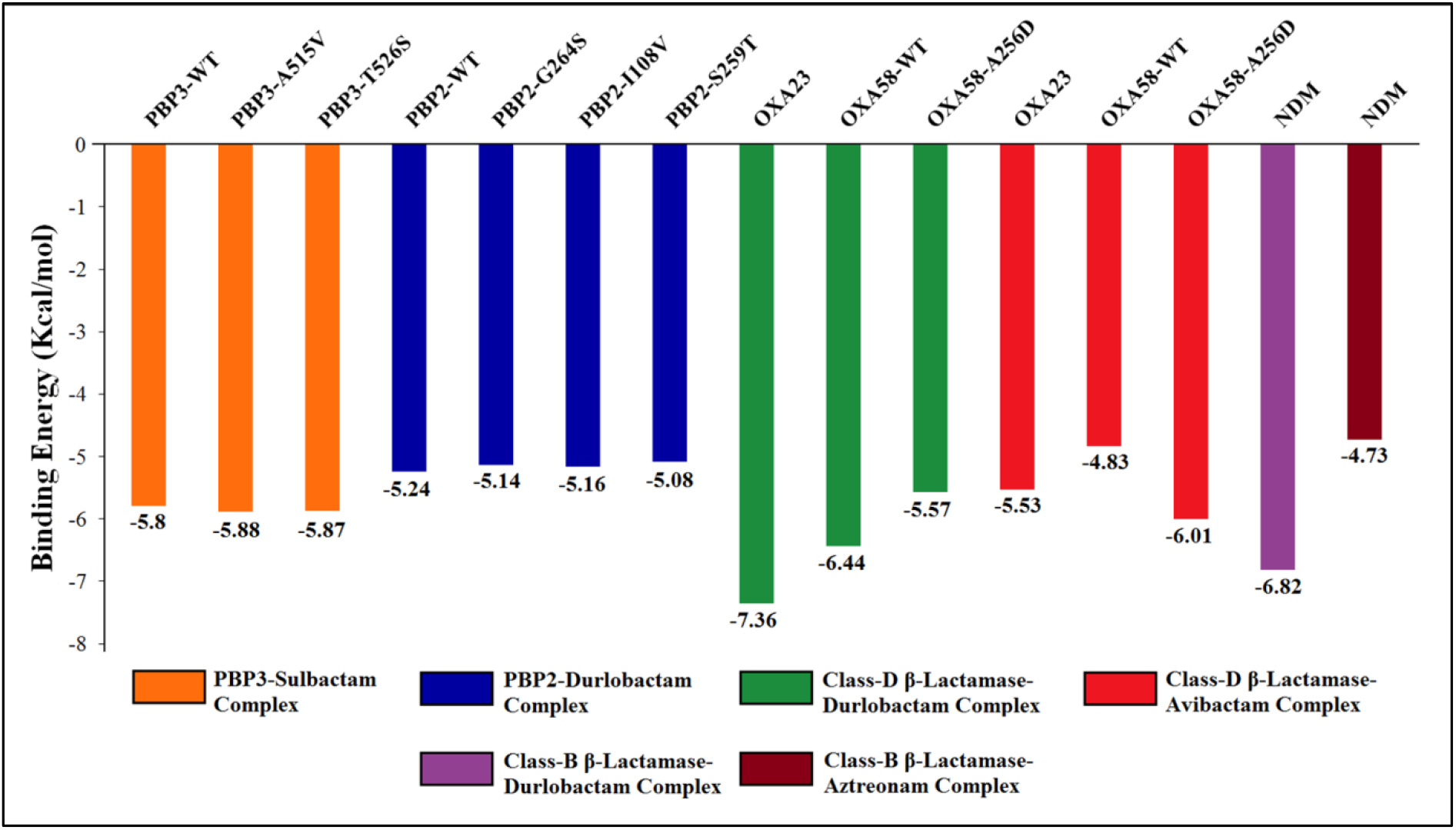
Binding Energies of Docked Complexes

There were no variations among the backbone-dynamics of STMK (average S^2^=0.71) and SSN (average S^2^=0.77) motifs of PBP3_WT_ and PBP3_MTs_ [**Supplementary File S2(B)**]. PBP3_T256S_ (average S^2^=0.65) and PBP3_A515V_ (average S^2^=0.7) lied within and in close vicinity of the KTGT motif respectively, resulting in flexible protein-backbone dynamics. However, there were no significant differences in BEs (−5.85±0.04Kcal/mol) and inhibition constants (IC) (51.36±3.7µM) of PBP3-SUL complexes. The flexible S336 residue (S^2^=0.71) facilitates in motif deprotonation followed by a nucleophilic attack through flexible K339 residue (S^2^=0.75) on the amide carbonyl-carbon of the βL-ring forming tetrahedral intermediate. Flexible residues of STMK and KTGT motifs stabilised the tetrahedral intermediate by stable H-bonds with the carbonyl O-atom of βL-ring inside the oxyanion hole [**Figure 5(A-C)**]. Finally, as tetrahedral intermediate collapses, the N-atom of βL-ring is expelled, forming a stable acyl-enzyme intermediate, thereby assuring proper inhibition of the target protein upon protonation by S390 residue (SSN motif) [King et al., 2017].

**Figure 5:**
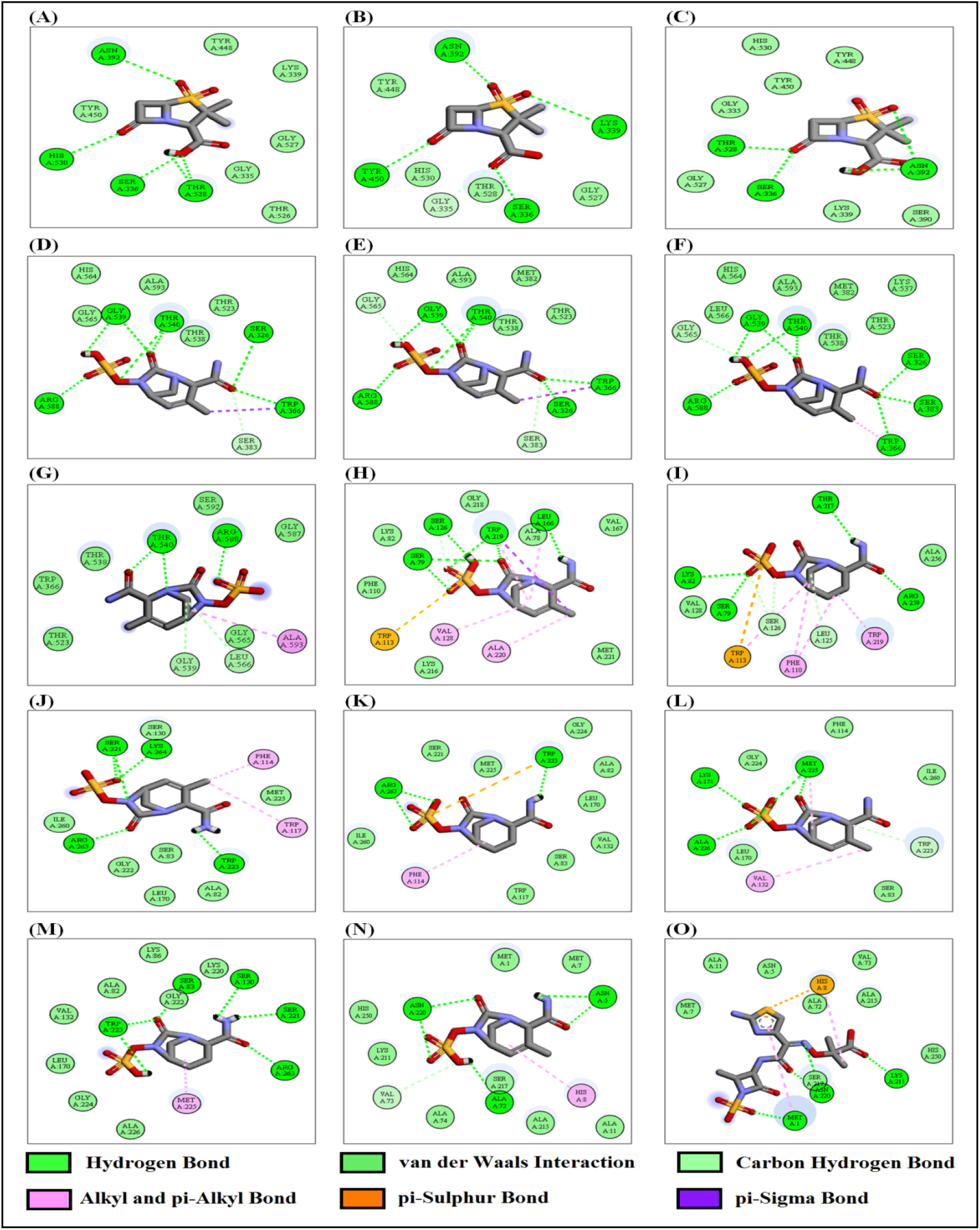
Docked poses of different protein-ligand complexes: **(A)** PBP3_WT_-SUL; **(B)** PBP3_T526S_-SUL; **(C)** PBP3_A515V_-SUL; **(D)** PBP2_WT_-DUR; **(E)** PBP2_G264S_-DUR; **(F)** PBP2_I108V_-DUR; **(G)** PBP2_S259T_-DUR; **(H)** OXA23-DUR; **(I)** OXA23-AVI; **(J)** OXA58_WT_-DUR; **(K)** OXA58_WT_-AVI; **(L)** OXA58_A256D_-DUR; **(M)** OXA58_A256D_-AVI; **(N)** NDM-DUR; **(O)** NDM-AZT

In PBP2, though individual residue-level analysis of nsSNPs revealed minimal fluctuations altering backbone flexibilities, no changes were noted in S^2^-values in any of the PBP2 active-site motifs [**Supplementary File S2(B)**]. The observation might be a crucial factor determining similar BEs (−5.16±0.66Kcal/mol) of PBP2-DUR complexes. An approximate 20% increase in IC was recorded for PBP2_MTs_ when compared with PBP2_WT._ PBP2-DUR interaction incorporated active S326, K329 residues of dynamic STIK (average S^2^=0.75) and G539, T540 residue of KTGT (average S^2^=0.72) motifs forming stable H-bond interactions with the carbonyl O-atom of DBO ring inside the oxyanion hole [**Figure 5(D-G)**]. OXA23-DUR (BE: −7.36Kcal/mol; IC: 4.01µM) displayed better interaction profile than OXA23-AVI (BE: −5.53Kcal/mol). The carbonyl O-atom of the DBO ring in DUR was positioned inside the oxyanion hole, forming a stable H-bond with active S79 of STFK motif, which was not observed with the carbonyl O-atom of AVI. The OXA23-DUR complex was further stabilised with F110 and M221, forming stable vdW interactions from either side of the active-site groove. The aromatic ∏-ring of W113 formed ∏-S interaction with S-atom of DUR while W219 formed ∏-σ interaction providing additional stability to the OXA23-DUR complex [**Figure 5(H-I)**]. Similarly, OXA58_WT_-DUR displayed better interacting profiles (BE: −6.44Kcal/mol; IC: 19.07µM) than OXA58_WT_-AVI (BE: −4.83Kcal/mol). The complex was further stabilised by vdW interaction with active S83, M225 and ∏-alkyl interaction with aromatic F114 residue.

On the contrary, OXA58_A256D_-AVI possessed a slightly better docked profile (BE: −6.01Kcal/mol) than OXA58_A256D_-DUR (−5.57 Kcal/mol). Analysing the docked poses of OXA58_A256D_-AVI (6 H-bonds) revealed stable H-bond interaction inside oxyanion hole with carbonyl O-atom of the DBO ring in AVI with the active S83. The MT-complex was further stabilised with ∏-alkyl interaction of the DBO ring with M225 residue. The OXA58_A256D_-DUR complex was stabilised by 4 H-bonds and vdW interactions with active S83 and F114 residues. Mutation in OXA58 is extremely rare and therefore, though AVI has shown better profiles than DUR, the efficacy of DUR cannot be ruled out as BEs and interactions of OXA58_A256D_-DUR complex was relatively stable [**Figure 5(J-M)**]. Molecular docking analysis revealed better docking profiles for NDM-DUR (BE: −6.82Kcal/mol; IC: 10.01µM) than NDM-AZT (−4.73 Kcal/mol). Though both the ligands bound to the active N220 residue, the carbonyl O-atom of the DBO ring in DUR was positioned in the oxyanion hole thereby ensuring better stability of the NDM-DUR complex [**Figure 5(N-O)**]. The binding parameters of SUL-PBP3 and DUR with β-lactamases and PBP2 displayed stable interaction profiles when compared to their classical drugs AVI and AZT. The present study thus established the potency of SUL-DUR combination which was effective against both WT and MT strains.

## CONCLUSION

Nosocomial strains of *A. baumannii* have created havoc in recent times, making them extremely difficult to treat. The AST profiles have demonstrated alarming XDR profiles of the bacteria. The CR*Ab* strains prompted us to shift to newer ΒL-ΒLI combinatorial therapy to combat the virulence caused by the strain. WGS analysis and gel-electrophoresis revealed the presence of several Class-B and D β-lactamases resulting in SUL resistance in 100% of the studied isolates. DUR displayed stable interaction profiles with low ICs in inhibiting β-lactamases which further facilitated effective inhibition of PBP2. The current study also hypothesises that DUR might be highly effective in treating dual carbapenemase and strains harbouring NDM alone. The treatment for OXA58_A256D_ might prompt administration of DUR at a higher concentration to achieve effective inhibition, although encouraging further *in-vivo* validations. Combination therapy of SUL-DUR could thereby restore SUL activity, which has shown significant binding affinities towards both PBP3_WT_ and PBP3_MTs_. The overall study has displayed substantial evidences in establishing the efficacy of novel βL-βLI combination, to curb the pathogenicity of CR*Ab* strains.

## CONFLICT OF INTEREST

The authors declare that there is no conflict of interests

## FUNDING

This research was funded by the Indian Council of Medical Research (ICMR), Govt. of India, through the research grant **IRIS-ID: 2019-0810**.

## ACKNOWLEDGEMENT

The authors would like to thank the management of the Department of Clinical Microbiology, CMC-Vellore and School of Bio-Sciences and Technology, VIT-Vellore, for providing the necessary facilities to carry out this research work. AN sincerely thank ICMR for his research fellowship. The authors would also like to acknowledge Mr Soumya Basu, VIT-Vellore, for his intellectual inputs while writing the manuscript and Ms Monisha Priya, CMC-Vellore, for helping us in retrieving the target protein sequences from WGS data.

## AUTHOR CONTRIBUTIONS

**AN:** Data Curation, *In-silico* methods and analysis, Writing-original draft;

**SV:** *In-vitro* methods-WGS analysis, Writing-original draft;

**BL:** Manuscript correction, supervision;

**BAS** and **SC:** Isolate revival, PCR, *In-vitro* MIC, AST;

**SR:** Validation of methodology, software and manuscript review;

**AA** and **BV:** Conceptualisation, Project Administration, Funding Acquisition, Manuscript review

## Notes

### Competing Interest Statement

The authors have declared no competing interest.

